# Toolbox for fluorescent labelling of *Pseudomonas aeruginosa* across scales: from single cells to bacterial communities and host infection models

**DOI:** 10.64898/2026.08.21.746161

**Authors:** Mattéo Gérard, Charlène Cornilleau, Vinciane Saint-Criq, Merve Nur Tunç, Maxime Deforet, Romain Briandet, Steven L. Porter, Rut Carballido-López

## Abstract

Fluorescence microscopy is central to the study of bacterial cell biology, multicellular behaviours, and host–pathogen interactions. Bright, robust and photostable labelling is required for bacterial identification, sorting and quantitative analysis, driving continuous development of state-of-the-art labelling tools. Here, we developed a multi-color fluorescent cell labelling toolkit for Gram-negative bacteria carrying the *attTn7* site, using the opportunistic human pathogen *Pseudomonas aeruginosa* as a model. Cell labelling is achieved by constitutive chromosomal expression of genes encoding a choice of four novel fluorescent proteins, mNeonGreen, mJuniper, mLychee and mScarlet-I3, codon-optimised for *P. aeruginosa*. These reporters provide bright, stable fluorescence with minimal photobleaching and excellent spectral separation during long-term imaging of single cells, macrocolonies and biofilms. Chromosomal expression of mNeonGreen yielded brighter and more homogeneous labelling than expression of the same construct from a plasmid. Interestingly, multi-color labelling of macrocolonies using our constructs allowed us to uncover and monitor a reversal of cell migration within motile colonies, thus highlighting the novel dynamics of motile communities. Finally, we demonstrate their applicability in biologically relevant host-pathogen contexts by imaging both live and fixed *P. aeruginosa-* infected human airway epithelial cells. This versatile cell labelling platform enables reliable bacterial identification, segmentation, tracking, and quantitative fluorescence imaging across spatial and temporal scales, and is readily adaptable to most other Gram-negative bacteria as the *attTn7* integration site is well conserved.

## INTRODUCTION

Fluorescence microscopy is an essential tool in bacterial cell biology, enabling the investigation of cellular organisation, multicellular behaviours and host-pathogen interactions. Its power to resolve single-cell dynamics, from tracking of individual molecules to imaging structured communities, has transformed our understanding of how bacteria grow, divide, adapt, evolve, and cause disease. Reliable cell labelling requires bright, photostable and spectrally versatile fluorescent markers to ensure robust cell detection, segmentation and tracking, and to enable quantitative analysis of cellular features across spatial and temporal scales with minimal perturbation of cell physiology.

The most widely used methods to fluorescently label live bacteria include chemical staining and genetically-encoded expression of fluorescent proteins (FP). Chemical dyes allow rapid labelling of specific cellular structures: DNA can be selectively stained using DNA-intercalating agents dyes such as DAPI and SYBR (1–6) , membranes by lipophilic dyes such as NileRed and FM, and the bacterial cell wall can be stained using fluorescently labelled lectins or analogues of D-amino acids (6–11). However, chemical dyes are generally not species-specific, which limits their applicability to complex samples such as multispecies communities or environmental samples. In host-pathogen infection models, membrane dyes and DNA intercalants often co-stain host cell membranes and nuclei, making unambiguous identification of bacteria within infected eukaryotic cells difficult or impossible. Furthermore, lipophilic dyes diffuse between membranes, and all chemical labels are progressively diluted with each round of bacterial division, thus compromising quantitative analyses and precluding long-term imaging experiments.

FPs overcome many of these limitations and remain the labelling method of choice for live-cell imaging of genetically-tractable bacteria. FPs are genetically encoded, non-toxic, and applicable across diverse growth conditions without the permeability and toxicity challenges associated with organic dyes. Early generations of FPs displayed slow chromophore maturation, low brightness and high susceptibility to photobleaching, which restricted their applicability in experiments requiring intense or prolonged illumination. However, engineering efforts have produced a continuously expanding palette of FP variants with substantially improved brightness, maturation kinetics and photostability (12), which now span the visible spectrum, enabling multi-color labelling of samples. The genes encoding FPs can be chromosomally expressed or carried on plasmids. Plasmid-based expression systems are easier to construct and to deliver, and can produce higher fluorescence signals through plasmid copy number amplification. However, cell-to-cell variation in copy number produces population level heterogeneity in fluorescence, complicating quantitative analysis. Furthermore, plasmid maintenance often requires continuous antibiotic selection (13), which can affect bacterial fitness, gene expression and the very physiological processes under investigation (14). In the absence of selection, plasmids can be rapidly lost, rendering quantitative fluorescence measurements unreliable over extended experiments or infection studies. Chromosomal insertion overcomes these limitations by ensuring single-copy, uniform expression of the FP across the entire population, irrespective of growth conditions and without requiring antibiotic selection.

The opportunistic Gram-negative human pathogen *Pseudomonas aeruginosa* is responsible for a range of severe acute and chronic infections, particularly in immunocompromised and cystic fibrosis patients(15–17). It constitutes a major health concern because of its high intrinsic antibiotic tolerance, combined with multiple acquired resistances(18). The number of multidrug-resistant, extensively drug-resistant and pandrug-resistant strains is increasing(19), leading the World Health Organization to classif*y P. aeruginosa* among the critical priority pathogens for which research and development of new antimicrobial compounds is urgently needed (20). Beyond its resistance, *P. aeruginosa* is also notorious for its wide variety of virulence factors and its capacity to form robust biofilms, which make it a privileged Gram-negative model for biofilm and virulence studies (18). Fluorescence-based investigation is therefore central to *P. aeruginosa* research and requires robust labelling tools that remain reliable across timescales, from minutes to days, and across spatial scales, from single cells to intact biofilms. Yet, fluorescence-based approaches and FP labelling tools for *P. aeruginosa* remain underdeveloped relative to other model bacteria such as *Escherichia coli* or *Bacillus subtilis*, or to other clinically relevant pathogens such as *Staphylococcus aureus or Streptococcus pneumoniae*.

In *P. aeruginosa*, FP imaging still relies predominantly on plasmid-borne expression(21), using classical green fluorescent protein (GFP) derivatives(22–24) and early-generation red FPs such as mCherry(23) and mScarlet(25), which have well-documented limitations in brightness and/or maturation time (12). The mini-CTX integrative system, derived from the filamentous bacteriophage CTX, has been widely used to achieve stable single-copy chromosomal insertion of genetic constructs *in P. aeruginosa* and related Gram-negative bacteria such as *Pseudomonas putida*, *Burkholderia spp*., and *Yersinia pestis*(26). It exploits site-specific recombination at the *attB* site mediated by the phage integrase, resulting in a single-copy insertion of the cargo at the conserved locus. Integration is selected using an antibiotic resistance marker, which can subsequently be excised via Flp–FRT recombination to yield a marker-free insertion. CTX-based vectors have been widely used in *P. aeruginosa*, notably for promoter–reporter fusions and genetic complementation (27); but chromosomal integration is relatively inefficient, occurring at frequencies of only 10⁻⁸ to 10⁻⁷. In this context, the mini-Tn7 transposon system offers a robust alternative. It uses the phage Tn7 transposition machinery to mediate single-copy insertion at the conserved *attTn7* site, located immediately downstream of the conserved *glmS* gene in at least 20 Gram-negative species (26,28,29). Because *glmS* is neutral with respect to cell physiology and integration frequencies are approximately 10⁴-10⁵-fold higher than those for mini-CTX, the mini-Tn7 system provides a more efficient and broadly applicable strategy for stable chromosomal integration in Gram-negative bacteria (23,30–32). Following antibiotic selection, the selection marker can be excised by Flp–FRT recombination, yielding a stable, marker-free chromosomal insertion.

In this work, we developed a multi-color fluorescent labelling toolkit based on stable chromosomal expression of four recent generation FPs, codon-optimised for *P. aeruginosa* to maximise translation efficiency. Using the mini-Tn7 integration system(29), we generated chromosomally integrated single-copy chromosomal constructs constitutively expressing the yellow-green mNeonGreen (33), the cyan-blue mJuniper and the orange-red mLychee (12) and mScarlet-I3 (34), all of which offer improved brightness and photostability compared with previous FP generations while collectively covering three spectrally separable imaging channels. We validated our toolkit across a broad range of applications, demonstrating compatibility with single and multi-color time-lapse imaging of single-cells, structured communities (macro-colonies and biofilms) and cell infection models, as well as with flow cytometry. By enabling robust, multi-color quantitative cell labelling across spatio-temporal scales and imaging modalities, this toolkit provides a versatile platform for fluorescence-based investigation of *P. aeruginosa* biology and is directly transferable to any species harbouring the conserved *attTn7* site. Here, we introduce a chromosomally integrated fluorescent toolbox that enables quantitative imaging across spatial scales. We demonstrate its power by uncovering previously unrecognized population dynamics during swarming.

## RESULTS

### Construction of multi-color fluorescent reporter strains of *P. aeruginosa*

We developed four modular integrative vectors using the pUC18T-mini-Tn7T plasmid(29) as the backbone. We selected four recent-generation monomeric FPs with enhanced brightness, photostability and maturation kinetics suitable for multi-color analysis: mNeonGreen (excited by blue light, laser line 488 nm), mScarlet-I3 and mLychee (excited by green light, laser line 561 nm) and mJuniper (excited by blue light, laser line 405/440 nm) (Fig. 1A-B). mNeonGreen is a robust green FP twice as bright as enhanced GFP (EGFP) that has already proven its efficiency in *Pseudomonas*(35). mScarlet-I3 is a last-generation fluorophore, reported to be the brightest red FP currently available, with a faster maturation time (21.5 min)(12) than widely used red FPs such as mCherry and mApple(12). mLychee is a novel red FP that is less bright and slower maturing (36.4 min) than mScarlet-I3 but reported to be more photostable(12). Knowing these FPs fundamental characteristics, mScarlet-I3 is theoretically better suited for endpoint imaging with no repeated illumination due to its higher photobleaching rate compared to mLychee that is less bright but more photostable, making it better for live imaging. Finally, mJuniper is reported to be among the best available blue-FP, with very fast maturation (1.7 min), low photobleaching and high brightness compared to other blue FPs such as mTurquoise2(12). The last step was to select a strong constitutive promoter, for chromosomal (single-copy) FP expression in *P. aeruginosa*. Promoter strength was the main criterion as the construct was going to be inserted as a monocopy rather than on a multicopy plasmid, thereby limiting the protein expression. The P*_tac_* synthetic hybrid promoter, derived from *E. coli* and P*_sbA_* promoters from *Amaranthus hybridus,* shows very strong expression in Gram-negative bacteria but has been shown to affect growth in *Pseudomonas fluorescens*(35). We therefore chose the P*_c_* promoter from a Class III integron (36), which is also commonly used in Gram-negative bacteria, and has strong activity (35,36).

**Fig. 1.**
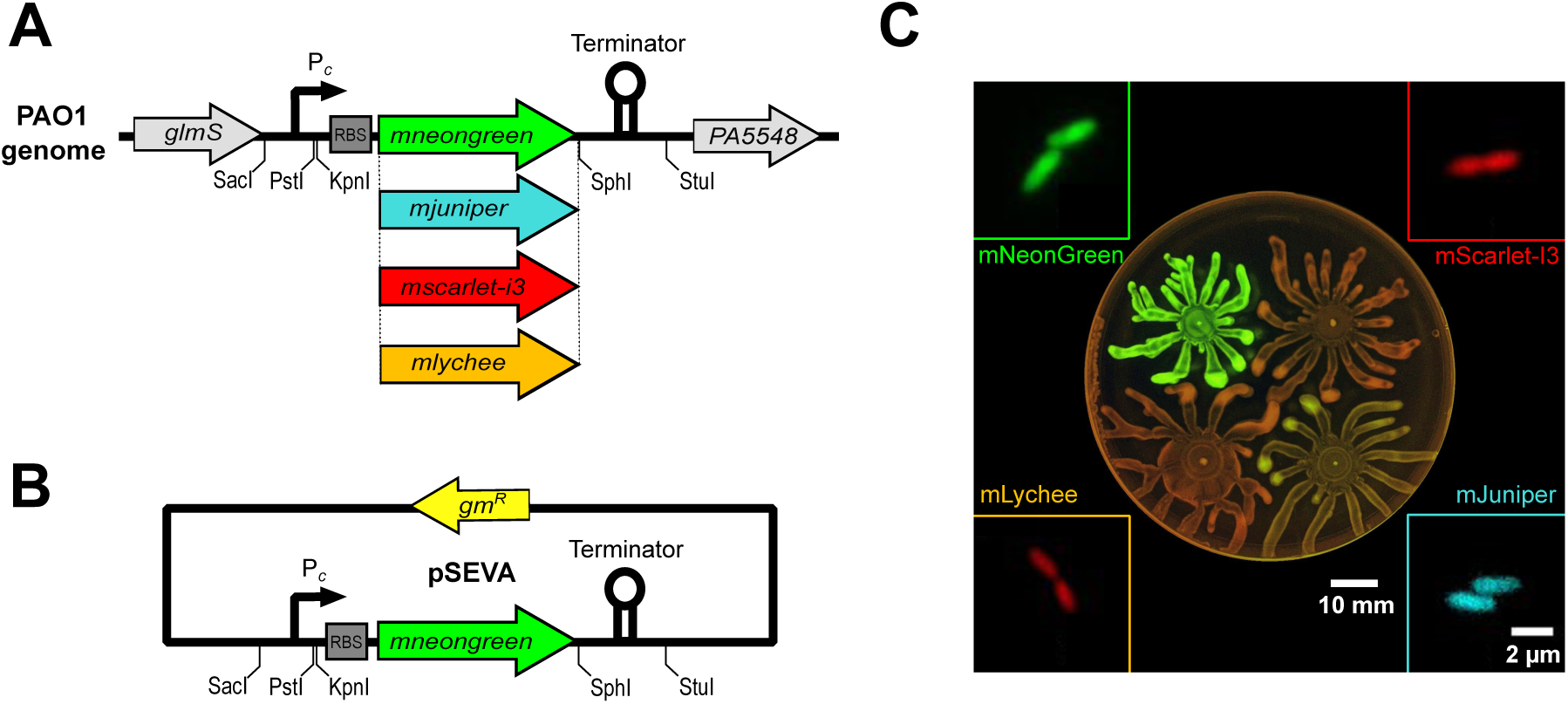
Genetic constructs and fluorophores presented in this study. **A, B,** Schematic of the five genetic constructs. Constructs are either chromosomally-inserted in the *P. aeruginosa* genome (A) or plasmid-borne (pSEVA) (B). Each chromosomal construct comprises the P***_c_*** promoter fused to the gene encoding one of four fluorescent proteins (mNeonGreen, mJuniper, mScarlet-I3, or mLychee), a Tn7 RBS, and the LUZ7 LUZT50 bidirectional terminator, flanked by unique restriction sites (SacI, PstI, KpnI, SphI, StuI) to allow modular exchange of each element. The pSEVA-derived plamid carries the same mNeonGreen construct. FP coding sequences were codon-optimised for *P. aeruginosa.* **C,** Representative colonies under UV illumination and single-cells imaged by conventional epifluorescence microscopy of strains P***_c_***-*mjuniper*, P***_c_***-*mneongreen*, P***_c_***-*mscarlet-i3*, P***_c_***-*mlychee*, chromosomally expressing the FPs after excision of gmR gene. Scale bars, 2 µm (for all cells) and 10 mm (colonies).

The pUC18T-mini-Tn7T-Gm plasmids carries the P*_c_* promoter (35) and the sequences of the genes encoding the four FPs codon-optimised for optimal expression in *P. aeruginosa*. The Tn7 major capsid protein ribosome-binding site (RBS)(35) was added upstream of the FP coding-sequence, and the LUZT50 bidirectional transcription terminator from *Pseudomonas* infecting phage LUZ7(37) was added downstream of the stop codon. Fragments were designed for straightforward insertion into pUC18T-mini-Tn7T using restriction enzymes and overlap-extension PCR. The resulting suicide vectors are pUC18T-mini-Tn7T-Gm-P*_c_*-mNeonGreenPA, pUC18T-mini-Tn7T-Gm-P*_c_*-mScarlet-I3PA, pUC18T-mini-Tn7T-Gm-P*_c_*-mLycheePA and pUC18T-mini-Tn7T-Gm-P*_c_*-mJuniperPA Each construct flanked by the Tn7 sites was integrated as a single copy at the *attTn7* site of the *P. aeruginosa* PAO1 genome using the Tn7 machinery, thus generating the 4 intermediate strains that contain each fluorescent construct and gentamicin marker. The gentamicin resistance gene was next excised using the Flp-FRT system of the pFLP3 plasmid, generating the four strains PAO1 *attTn7*::(P*_c_*-*mneongreen*) (pRCL240), PAO1 *attTn7*::(P*_c_*-*mscarlet-i3*) (pRCL236), PAO1 *attTn7*::(P*_c_*-*mlychee*) (pRCL244), and PAO1 *attTn7*::(P*_c_*-*mjuniper*) (pRCL248).

To compare chromosomal and plasmid-based FP expression, the P*_c_*-*mneongreen* construct was cloned in parallel into the pSEVA627M backbone(38). The resulting plasmid, pSEVA-Gm-P*_c_*-mNeonGreen (Fig. 1A) was electroporated into *P. aeruginosa,* thus generating strain PAO1 pSEVA-P*_c_*-*mneongreen* (pRCL251). The replication origin for this plasmid is derived from the broad-host-range RK2(39), which enables stable replication in several Gram-negative bacteria, including *Pseudomonas* species. RK2-based replicons are maintained at relatively low copy number, typically around 4-7 copies per cell in *E. coli* and approximately 3 copies per cell in *P. aeruginosa*(39).

All plasmids were designed for both stable expression in *P. aeruginosa* and straightforward modularity. Each element of the constructs is flanked by unique restriction enzymes sites (Fig. 1A), allowing the promoter, FP, or transcription terminator to be exchanged for the desired sequence.

All strains displayed growth comparable to that of the parental PAO1 wild-type strain, either after or before excision of the gentamicin cassette, and a bright FP signal at both the single cell level and in colonies (Fig. 1C).

### Suitability of the fluorescent reporter strains for long-term single-cell imaging

We first assessed the fluorescence intensity and photostability of the constructs at the single-cell level. Cells were imaged by epifluorescence microscopy in 2-hour live-cell time-lapse experiments using low laser power and a 5-minute acquisition interval (Fig. 2A). We imaged the four strains carrying chromosomally integrated constructs, as well as the strain expressing the mNeonGreen construct from a plasmid. The plasmid carrying strain was exposed to gentamicin throughout the experiment to maintain plasmid stability. Each strain displayed homogenous, bright cytoplasmic fluorescence signal (Fig. 2A). Quantitative analysis of fluorescence intensity over the 2-h time-lapses showed that all four FPs retained stable fluorescence (Fig. 2B-D). Chromosomally expressed mNeonGreen remained highly photostable throughout the first hour of imaging, showing only modest photobleaching thereafter. Surprisingly, it was two-fold brighter than when expressed from the pSEVA plasmid, which in contrast underwent most of its photobleaching during the first hour before reaching a stable plateau (green and purple, respectively in Fig. 2B). This unexpected observation that the same P*_c_*-*mneongreen* construct displayed a substantially higher fluorescence when expressed from the chromosome than from the pSEVA-derived plasmid (Fig 2A, B), prompted us to compare the two strains in more detail. Flow cytometry analysis of liquid cultures showed a unimodal fluorescence distribution for both strains, confirming homogeneous protein expression at the population level, but plasmid-based expression gave a slightly broader fluorescence distribution, consistent with fluctuations in RK2-based plasmid copy number and/or differences in plasmid segregation between cells. By flow cytometry the difference in signal intensity was more pronounced than in epifluorescence, with chromosomally tagged P*_c_*-*mneongreen* displaying approximatively ten times higher brightness per cell than the plasmid-based construct. The mini-Tn7 system provides single-copy chromosomal insertions, whereas pSEVA-derived plasmids are maintained at ∼ 3 copies per cell in *P. aeruginosa*(39). Despite this difference in copy number, the P*_c_*-*mneongreen* construct showed consistent higher brightness when chromosomally-expressed at the single-cell level. Plasmid-mediated expression also showed faster photobleaching and greater cell-to-cell signal heterogeneity (Fig. 2B).

**Fig. 2.**
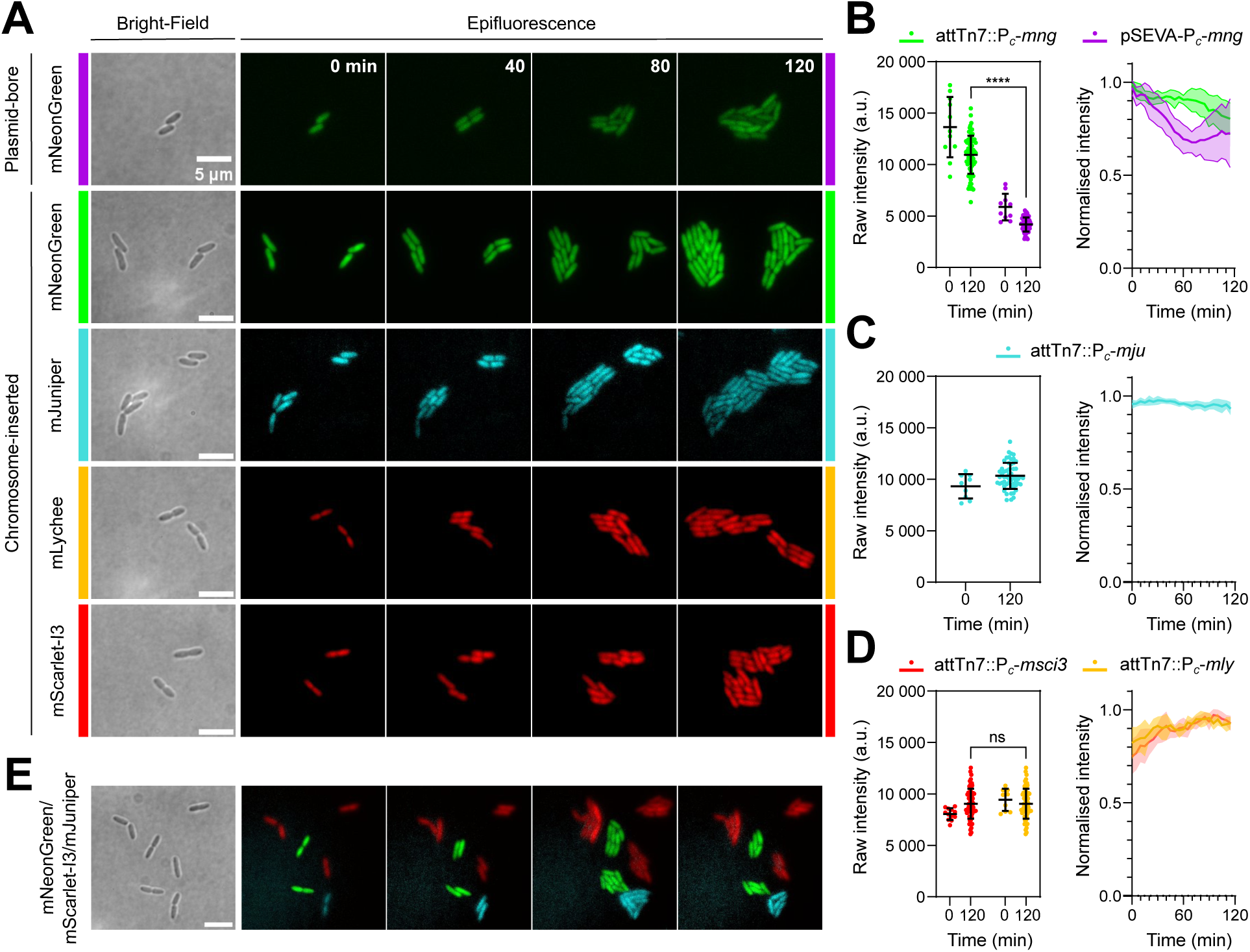
The four chromosomally-inserted constructs show high brightness and photostability, and are suited for multi-color imaging at the single-cell level. Violet, pSEVA-P*_c_*-*mneongreen*; green, chromosome-inserted P*_c_*-*mneongreen*; blue, P*_c_*-*mjuniper*; yellow, P*_c_*-*mlychee* and red, P*_c_*-*mscarlet-i3*. **A,** Epifluorescence time-lapses in LB media. All strains were imaged individually for 2 h with a 5 min acquisition interval, and 100 ms exposure time. Representative bright-field at time 0 and epifluorescence images at 0, 40, 80, and 120 min are shown. Scale bars, 5 µm. **B-D,** Brightness and photostability of the constructs over the full-time lapse. In the scatter plots, the main bar indicates the mean and error bars the standard deviation. Line plots represent normalised to maximum signal intensities over the 2-h time-lapse (mean ± SD), total n>50 cells in 3 biological replicates. (B) Plasmid-borne and chromosomally inserted P*_c_*-*mneongreen*; ****, P-value < 0.0001 (nonparametric two-sided Mann–Whitney test). (C) Chromosomally inserted P*_c_*-*mjuniper*. (D) Chromosomally inserted red FP fusions P*_c_*-*mscarlet-i3* and P*_c_*-*mlychee*; ns, P > 0.05 (nonparametric two-sided Mann– Whitney test). **E,** Tricolour imaging of the of P*_c_*-*mneongreen*, P*_c_*-*mjuniper* and P_c_*-mscarlet-i3* mixed at a 1:1:1 ratio. Scale bar, 5 µm.

While all chromosome-inserted fluorophores displayed stable expression with greater than 75 % of the initial fluorescence remaining after the end of the 2 h time lapse, mJuniper was the most stable having close to 100% remaining. (Fig. 2C). Interestingly, the two red FPs displayed a similar intensity profile over time, with approximately 20 % rise in signal over the first 40 min, followed by a plateau (Fig. 2D). This behaviour is consistent with the slower maturation half time of these two proteins (mLychee, 36.4 min; mScarlet-I3, 21.5 min). We next used flow cytometry to compare the fluorescence intensity and distribution of the two red FP expressing strains, mScarlet-I3 and mLychee. Both showed similar peak intensity profiles, but mLychee displayed a left-skewed distribution towards lower-intensity signals compared to mScarlet-I3. The origin of this atypical distribution remains undetermined, yet it does not affect signal homogeneity in epifluorescence studies (Fig. 2D).

We also compared the strain carrying the chromosomal mNeonGreen construct with the intermediate strain carrying the same construct but before excision of the gentamicin-resistance cassette, whose presence showed no impact on growth rate. The two strains displayed comparable cell labelling over 2 h epifluorescence time-lapses, and flow cytometry revealed no significant difference in signal intensity. Since no gentamycin was added to the culture media, continuous presence of antibiotic is not required to maintain chromosomal constructs, resistance-cassette excision appears as an optional step.

Finally, we tested fluorophore compatibility for multi-color imaging by mixing three strains chromosomally expressing proteins with non-overlapping spectra, mNeonGreen, mJuniper and mScarlet-I3, at a 1:1:1 ratio (Fig. 2E). No bleed-through between channels was observed, and the signals from the three fluorophores were clearly distinguishable.

Taken together, these data suggest that our constructs are suitable for single-cell live single and multi-color imaging. mNeonGreen displays high brightness but bleaches slightly faster than the other fluorophores, although bleaching remains very low under our imaging conditions. mJuniper displayed the most stable signal over time, consistent with its very fast maturation. The two red FPs, mScarlet-I3 and mLychee, displayed high intensity signals suitable for long-term imaging but may be less suited for monitoring subtle or fast dynamics owing to their slow maturation times. Chromosome-based outperformed plasmid-based P*_c_*-*mneongreen* expression in signal intensity, stability and signal homogeneity at the single cell level. It is important to note that gentamycin was used only in the plasmid-based setup: continuous selection during imaging indicates that the observed difference is unlikely due to plasmid loss and more likely attributable to expression disparities between the single-copy chromosomal locus and the low-copy plasmid and/or to antibiotic effects: gentamicin targets ribosomes thereby slows down translation; even though plasmid carrying cells expresses gentamicin resistance gene, residual antibiotic effect can remain and affect mNeonGreen expression. This residual antibiotic activity could explain measured differences in signal intensities between chromosome and plasmid-based expression.

### Evaluation of cell labelling efficiency of the engineered constructs in biofilms

We next assessed cell labelling of the engineered strains within complex bacterial communities by imaging submerged biofilms using confocal laser scanning microscopy (CLSM) (Fig. 3A). The four chromosomally tagged strains were grown as submerged biofilms and imaged after 4, 12, or 24 hours to follow the characteristic stages of *P. aeruginosa* biofilm formation(40,41); mean biofilm thickness was quantified at each timepoint for each strain (Fig. 3B). After 4 h, scattered micro-colonies had formed and proliferated laterally across the surface, which was not yet fully covered. By 12 h, microcolonies had coalesced and biofilms had matured into a thin, structured layer of 20-40 µm thickness. All four strains displayed stable or slightly increasing thickness between 4 and 12 h, consistent with lateral proliferation; and by 24 h, they had matured into a thick, dense layer(40,41) with a 2 fold increase in thickness to 50-60 µm. Stable chromosomal expression ensured persistent fluorescence throughout biofilm maturation without the need for antibiotic selection, a key advantage for long-term imaging experiments. Further structural analyses were performed (Fig. 3C-D) by quantifying biofilm biovolume and surface roughness for each strain. Biovolume increased consistently from 4 to 24 h, reflecting progressive biofilm development and biomass accumulation (Fig. 3C). Concomitantly, biofilm roughness decreased over time, consistent with the formation of a dense and more homogeneous biofilm architecture as the biofilms matured (Fig. 3D). Performed quantifications also allowed for precise comparison of each strain at each timepoint. Statistical analyses were performed using the mNeonGreen strain as a reference: on the one hand, these revealed variability between strains for each extracted data (Thickness, Biovolume, Roughness) in maturing early biofilm stages (Fig. 3B-D) for all parameters, and does not seem to be specific of one single strain. On the other hand, all strain showed no statistically significant differences with reference for the three tested parameters in the 24 h mature biofilm. These data suggest that differences observed in early biofilm stages may be due to variability in biofilm structure during maturation, and that all strains enable precise biofilm quantification, as they perform equally well in stable, mature structures.

**Fig. 3.**
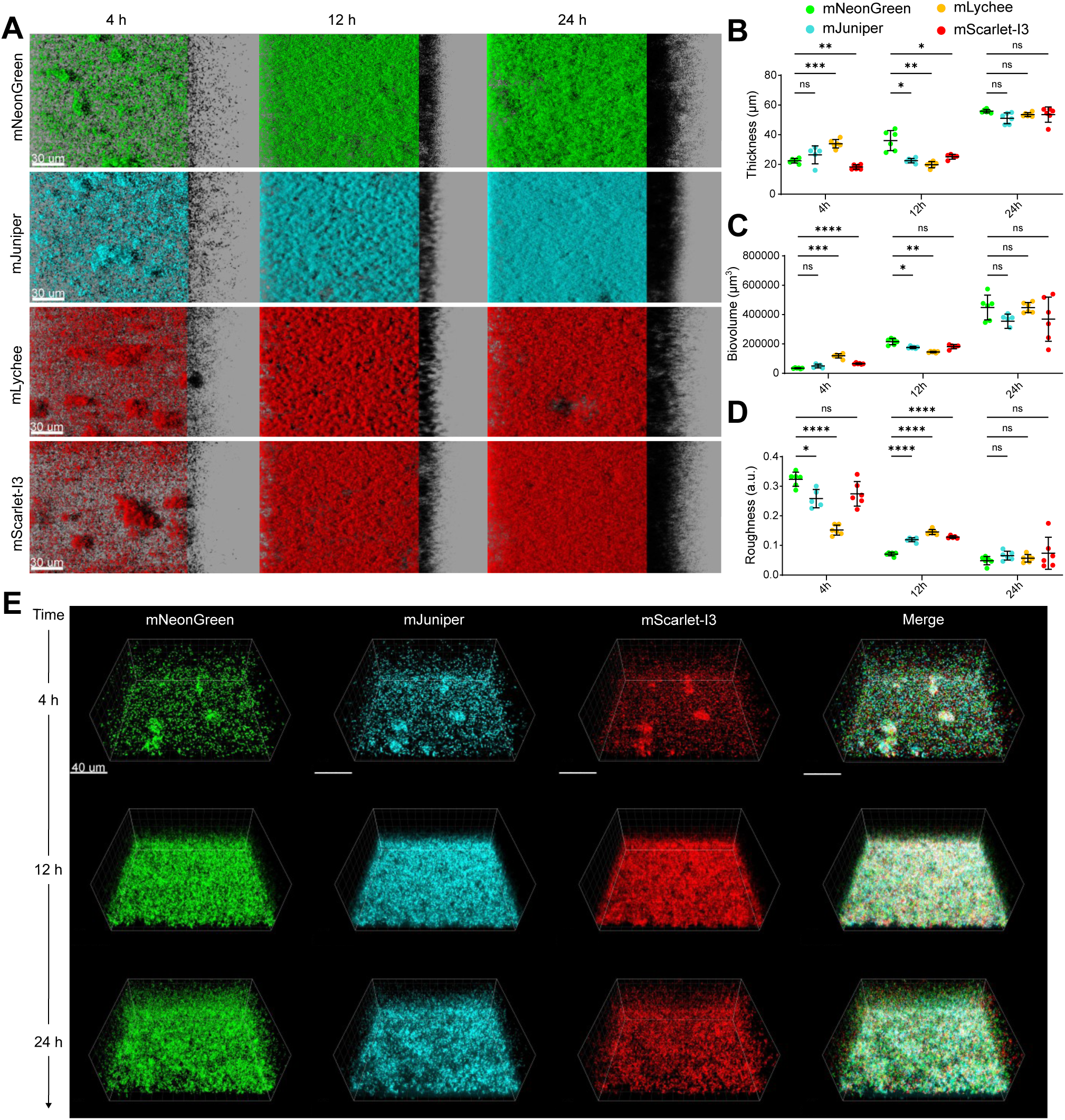
The four chromosome-inserted strains provide clear visualisation of biofilm architecture and development. **A**, Top view (xy) of confocal laser scanning microscopy (CLSM) images from 4, 12 and 24 h biofilms of the four strains indicated, carrying chromosome inserted tags. Side biofilm view is represented with a black shadow. Scale bars, 30 µm. **B-D,** Mean biofilm thickness (µm) (B) biovolume (µm**^3^**) (C) and roughness (a.u.) (D) based on the CLSM stacks of the time points shown in (A) Each dot represents a technical replicate (5 stacks per strain); the central bar represents the mean and error bars are standard deviation. ****, P < 0.0001; ***, 0.0001 < P < 0.001; **, 0.001 < P < 0.01; *, 0.01 < P < 0.05; ns, P > 0.05. (Tukey’s multiple comparisons test) **E,** Three-dimensional (xyz) CLSM images of 4, 12 and 24 h mixed biofilms composed of three strains (attTn7::P***_c_***-*mneongreen*, attTn7::P***_c_***-*mscarlet-i3* and attTn7::P***_c_***-*mjuniper*) mixed at a 1:1:1 ratio at the pre-adhesion stage. Each fluorescence channel is shown separately, and the merge combines the three channels. Scale bar, 40 µm.

Ultimately, all four constructs enabled monitoring of biofilm formation by *P. aeruginosa*, yielding a bright, evenly distributed signal that clearly reported biofilm architecture at each stage. No strain displayed fluorescence signal that would not be suited for biofilm imaging, nor channel-specific artefacts. Importantly, fluorescence remained homogeneous throughout the biofilm volume, with no detectable attenuation along the z-axis, enabling reliable quantitative analysis of the entire three-dimensional community. Quantitative readout of biofilms growth through mean thickness, biovolume and roughness quantification in CLSM stacks showed a similar evolution for all four strains, with a consistent biovolume increase over time, with the strongest growth occurring between 12 and 24 h as biofilms transition to their mature, thick-layer stage (Fig. 3B). Despite the statistically different growth in between strains observed in early biofilm stages, differences ultimately remained low and cannot be linked to defected growth of one of the strains. This is consistent with their shared genetic background and liquid cultures growth rates, thus indicating that all of the FPs are compatible with strong biofilm growth. In summary, quantifications trajectories mirrored the stages visualised in CLSM images, confirming that all reporters faithfully report biofilm formation over time. The combination of homogeneous fluorescence and stable reporter expression makes these strains particularly suitable for automated image segmentation and quantitative three-dimensional biofilm analyses.

We next investigated whether several chromosomally tagged strains could be simultaneously visualised to resolve distinct subpopulations within biofilms. The three strains constitutively expressing mNeonGreen, mJuniper and mScarlet-I3 were mixed at a 1:1:1 ratio at the pre-adhesion stage and grown together as a mixed biofilm; each strain grew evenly over time, following the same biofilm development pattern observed in single strain imaging (Fig. 3E). The three populations were clearly separated in their respective channels, with no detectable bleed-through (Fig. 3E), and the merge image showed that they intermixed evenly throughout the biofilm volume at all time points. Each strain co-developed with the same kinetics observed in the single-strain biofilms, from sparse surface-associated cells at 4 h to a thick, three-dimensional community by 24 h. Together, these data indicate that the three spectrally separated FPs support high-quality biofilm labelling and can be combined to track bacterial subpopulations in complex structured communities, a capability that could be extended to strains of different genotypes or to mixed-species consortia. Such stable multicolour labelling provides a quantitative framework for investigating spatial organization, mutant competition, cooperation and niche partitioning within mono-and multispecies biofilm communities.

Overall, each genetic construct supports high-quality biofilm labelling, from initial surface colonisation to the mature biofilm. No strain exhibited an altered biofilm phenotype thus confirming the absence or minimal impact of the constructs on biofilm fitness. These data establish our reporter set as a robust tool for long-term, single-and multi-strain biofilm imaging.

Lastly, we compared the plasmid and chromosome-based mNeonGreen expression strains as biofilms to investigate whether the single-cell differences were carried over to a community context. Biofilm architecture, thickness, roughness, and biovolume showed similar evolutionary patterns for both strains and showed statistically significant differences between them at each timepoint, indicating that both report biofilm development equally well. To assess whether labelling or antibiotic selection affected viability, biofilms were labelled with 1 µM propidium iodide (PI), which only enters cells with compromised membrane, thereby allowing evaluation of the proportion of dead and damaged cells. PI-positive cells were rare in young biofilms but became more frequent by 24 h, concentrated in the deeper layers, consistent with increased lethality in maturing *P. aeruginosa* biofilms (1,42). No extra death was observed in between conditions and biofilm structure remained similar, even though the plasmid-tagged strain was grown in media containing 50 µg/mL gentamicin. Chromosomal expression of the four FPs provides stable, homogeneous labelling of developing and mature biofilms without detectable fitness cost, enabling quantitative three-dimensional analyses of single-and mixed-population biofilms

### Dual-color labelling of swarming colonies reveals *P. aeruginosa* population dynamics

To move beyond validation and demonstrate biological utility of our toolkit, we tested our fluorescent strains in macroscopic swarming motility assays. Swarming motility is a major determinant of *P. aeruginosa* virulence(43) and has been the subject of extensive investigation. It is a flagellum-driven collective movement on semi-solid surfaces, characterized by the circular, dendritic expansion of bacterial colonies, and associated with rhamnolipids production, flagella expression and quorum sensing(44,45). It is considered clinically relevant because the semi-solid swarming surface mimics the thickened mucus of infected airways, and swarming cells exhibit increased antibiotic tolerance (46).

Macroscopic images of 24 hours swarming colonies were acquired under blue LED excitation and white-light illumination using a Reshape imaging system (Reshape Biotech, Denmark) (Fig. 4A). The strain chromosomally expressing mNeonGreen exhibited a typical swarming phenotype, forming radially expanding dendritic structures(45) with higher cell density at the tendril tips, consistent with previous reports(47) (Fig. 4A). Fluorescence imaging revealed a heterogeneous distribution of mNeonGreen-expressing cells across the colony, with uneven signal along the tendrils and increased signal at the colony periphery, particularly at the tendril tips (Fig. 4B). Quantification of fluorescence intensity across the tendrils branches and at their tips revealed an ∼ 3-fold higher fluorescence at the tips (Fig .4B), consistent with the increased cell density observed under white-light illumination. This tip enrichment is consistent with the leading edge of a swarm being the most densely packed, metabolically active zone, where actively dividing cells accumulate ahead of the following cell monolayer (48). We conclude that fluorescence intensity reflects cell density at the swarming front, although this relationship remains approximate due to potential variations in metabolic activity, oxygen availability or growth phase that may produce uneven signal distribution across the colony. Nevertheless, the overall fluorescence distribution closely tracks cell density making it a proxy for bacterial distribution within the swarm.

**Fig. 4.**
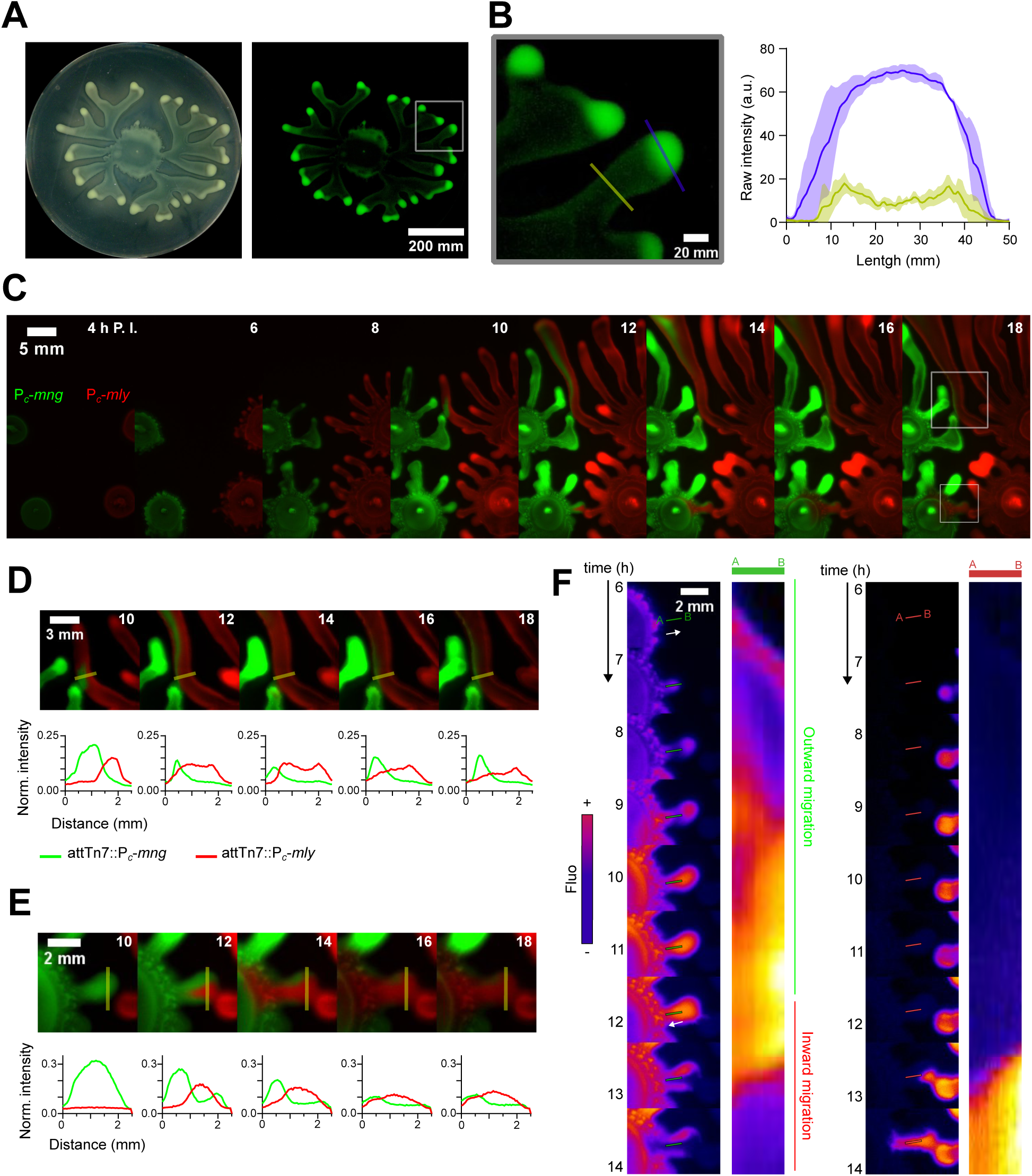
Multi-color reporter strains reveal swarming flow dynamics. **A**, 24 hours swarming colony of *P. aeruginosa* carrying attTn7::P*_c_*-*mneongreen* construct, imaged on a Reshape imaging system under white-light illumination (Left) and blue LED excitation (Right). **B,** mNeonGreen intensity quantification was performed in two distinct regions of the swarming colony: across the tendril shaft (yellow line) and at the tendril tip (swarming front, purple line), where cell accumulation is maximal. Fluorescence intensity was measured along 5 mm profiles from 4 tendrils of similar width. Lines show the mean and dashed areas the standard deviation. **C,** Dual-color swarming time-lapse of the P*_c_*-*mlychee* and P*_c_*-*mneongreen* strains grown in the same plate, from 4 to 18 h post-inoculation (P. I.). **D, E,** Mixing dynamics of two motile colonies (10-18 h P.I.). Fluorescence intensity was measured along 2.5 mm slices across the contact zone every 2 h (yellow lines); profiles below each frame show normalised mNeonGreen (green) and mLychee (red) intensity. **F,** Cell migration within P*_c_*-*mneongreen* colony. Heatmap reveals fluorescence intensity (yellow-high to purple-low) in both green and red channels. Kymographs shows intensity over the A to B line between 6 and 14 h P.I. for either green or red fluorescence, the white arrow indicates cell migration direction.

We next performed dual-color swarming fluorescence time-lapses of the strains carrying chromosome-inserted P*_c_*-*mneongreen* and P*_c_*-*mlychee* constructs inoculated in close contact and grown side-to-side on the same plate (Fig. 4C). Because the two strains have the same genetic background, this setup allows for the investigation of specific phenomena: how tendrils expand when neighbouring colonies approach, and how two genetically identical colonies mix. We observed the characteristic two-step swarming expansion of *P. aeruginosa*(49): after inoculation, each colony expanded as a circular, outward-spreading disc during the first 6 h, before branching began and tendrils formed around the colony edge (Fig. 4C).

By inoculating colonies in close contact, our experimental setup forces them to meet, allowing us to monitor how two swarming populations interact (50). We then investigated the mixing dynamics of motile *P. aeruginosa* subpopulations across 2.5 mm slices in the contact zone (Fig. 4D-E). This revealed two interesting population dynamics. First, tendrils expanding in similar directions occasionally can fuse (Fig. 4D). When the tip of the green tendril met the side of the red tendril, a fraction of the green population migrated towards the red tendril tip. Consistently, the intensity profiles across the fusion zone show the green signal progressively displaced along the red tendril (Fig. 4D). Second, cell migration inside the colony is highly dynamic (Fig. 4E). Between 4 and 8 h after inoculation, each colony expanded radially until two tendrils came into direct contact tip to tip. Unexpectedly, the colonies then merged but did not mix evenly; only the red strain (P*_c_*-*mlychee*) invaded the originally green colony (P*_c_*-*mneongreen*); the intensity profiles across the contact slice show the red signal rising and the green signal receding over time (Fig. 4E). This produced a retrograde movement inwards in the direction of the inoculation spot, while the red cells advanced inside the green colony until they reached the inoculation spot. Because movements within the colony are expected to expand outwards from the inoculation spot, this inward flow after tendril contact was unexpected.

To further characterize the inward migration dynamics, we generated kymographs of fluorescence intensity from the P*_c_*-*mneongreen* colony (Fig. 4F). Green fluorescence intensity was monitored along the cell flow axis over time. During the first 12 h, the kymograph revealed a clear outward expansion of the colony, with fluorescence intensity progressively increasing along the measured axis. After 12 h, however, fluorescence intensity began to decrease from the opposite end of the axis, providing direct evidence of a reversal in the direction of cell flow within the colony (Fig. 4F). Such large-scale rearrangement is consistent with hydrodynamic flows generated by swarming monolayers and may be related to processes involved in swarming regulation, such as local rhamnolipid gradients or quorum-sensing cues (51). These data indicate that cells remain highly motile and are carried along the entire length of the tendril, not only at the tendril tip, consistent with the idea that swarms are not simply expanding fronts but maintain internal circulation and cell exchange throughout the tendril (25,48)

Overall, our fluorescent tools allowed for robust discrimination of two isogenic populations at the macroscopic scale and revealed both anticipated behaviours (two-step expansion, tendril self-avoidance) and previously unrecognized internal migration-dynamics (asymmetric invasion and tendril fusion) within swarming colonies. Beyond validating our reporters, these observations show that constitutive multi-color chromosomal labelling is well suited to studying collective motion and population mixing in *P. aeruginosa* swarms, a system where most prior imaging has relied on single-color or transmitted-light readouts.

### Imaging of *P. aeruginosa* epithelial cell infection

Finally, we characterised the virulence of the engineered *P. aeruginosa* strains in infection models, which pose several imaging challenges. These include limited sensitivity in fluorescence detection, constraints on bacterial and host cells specific labelling, sample complexity, and the need to maintain physiological conditions during live imaging. To address these, we monitored infection dynamics using bronchial epithelial BEAS-2B cells.

We first performed a 10 h widefield time-lapse of epithelial cells infected with the chromosomally inserted P*_c_*-*mneongreen P. aeruginosa* strain (Fig. 5A). BEAS-2B mitochondria were labelled with MitoTracker Deep Red FM before being infected with a multiplicity of infection (MOI) of 15 and imaged for 10 h. In the non-infected control condition, cells stretched and adhered to the surface over time, reflecting cytoskeleton remodelling during adhesion (Fig. 5A, *Top panel*). In the infected condition, bacteria adhered to the surface of the epithelial cells, which rounded up (Fig. 5A, *Bottom panel*). Such rounding can be attributed to virulence factors that collapsed the actin cytoskeleton(52) or drove rapid membrane damage and cell lysis (53). Infection was performed in PBS so that bacteria could not grow using the culture media. Over time, bacteria lysed the host cells (as shown by loss of cell integrity and release of intracellular content) and proliferated on the released cell lysate or dead cells (Fig. 5A), consistent with the pore-forming and phospholipase activities that make *P. aeruginosa* acutely cytotoxic to airway epithelia (53). Bacterial fluorescence and the number of non-lysed epithelial cells were quantified at each timepoint (Fig. 5A-B). Cell lysis occurred between 2-and 3-h post-infection, and cell count reached zero by ∼3.5 h. Bacterial fluorescence rose as the host cells lysed, peaking around 7 h before declining over the remainder of the time-lapse (Fig. 5A-B). This is consistent with a switch from bacterial adhesion/intoxication to a replicative phase induced by the nutrients released from lysed host cells, until nutrient exhaustion and dispersal from the field of view. Throughout the 10 h time-lapse, the chromosome-based P*_c_*-*mneongreen* construct displayed high brightness and photostability, allowing precise localisation and quantification of bacteria during infection.

**Fig. 5.**
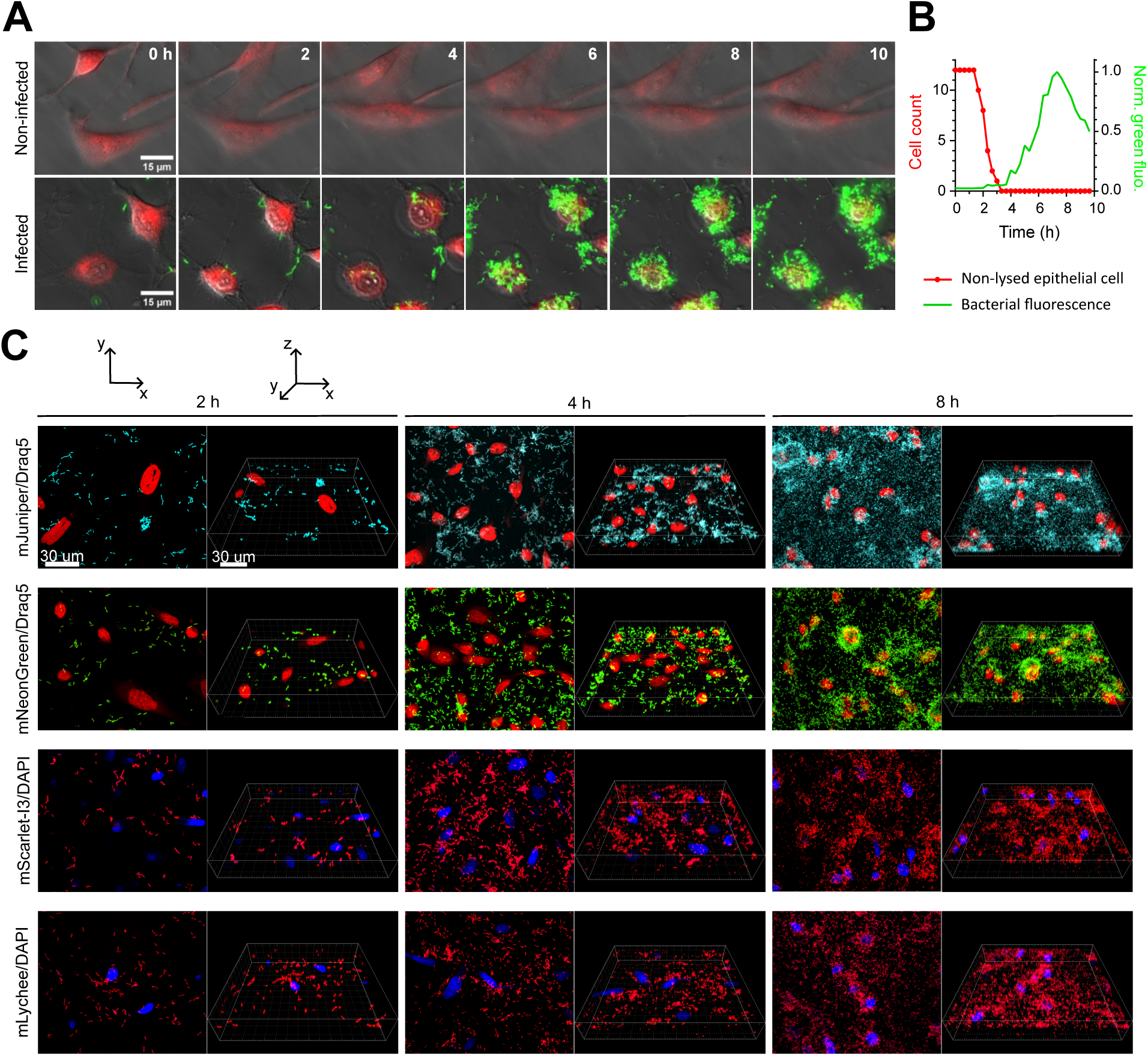
High precision bacterial tracking in a host-infection model. **A**, Widefield fluorescence timelapse of BEAS-2B bronchial epithelial cells infected with the attTn7::P*_c_*-*mneongreen* strain (green). Host mitochondria were labelled with 200 nM Mito Tracker Deep Red FM prior to infection. Cells were infected at a MOI of 15 and monitored for 10 hours. Uninfected (Top panel) and infected (Bottom panel) conditions are shown at the indicated times. **B,** Quantification of infection dynamics from (A). The green line represents the normalised bacterial fluorescence over the full camera field; the red line represents the non-lysed epithelial cells at each time point. **C,** CLSM imaging of fixed BEAS-2B cells infected with each of the four chromosomally tagged strain (green, P*_c_*-*mneongreen*; cyan, P*_c_*-*mjuniper*; red, P*_c_*-*mscarlet-i3* and P*_c_*-*mlychee*) after 2, 4, and 8 h of infection. Host nuclei were counterstained with 1 µM DAPI (mScarlet-I3, mLychee) or 3.75 µM Draq5 (mNeonGreen, mJuniper) prior to imaging. For each strain and timepoint, the left image is a top view (xy) of the full stack and the right image a three-dimensional (xyz) view of the same stack. Scale bars, 30 µm.

We then increased the imaging resolution using confocal microscopy, and extended the assay to all four chromosomally tagged strains. BEAS-2B epithelial cells were infected with each strain individually and imaged by CLSM on fixed samples after 2, 4 and 8 h of infection (Fig. 5C). Host nuclei were counterstained with Draq5 (for mJuniper and mNeonGreen) or DAPI (for mScarlet-I3 and mLychee). The four FPs displayed high brightness and stability at each post-infection time point, and in every case the FP signal was spectrally well separated from the counterstain, allowing bacteria and host nuclei to be resolved in distinct channels. Progressive bacterial growth across the epithelial layer was observed, with sparse, well-separated bacteria and intact nuclei at 2 h giving a dense bacterial coverage by 8 h (Fig. 5C). Together, these data demonstrate that all four reporters enable accurate three-dimensional bacterial localization and resolve individual bacteria against a labelled host background, establishing their suitability for infection imaging studies.

## DISCUSSION

In this work, we generated and validated a set of genetic constructs for efficient multiscale labelling of *P. aeruginosa*. These are single-copy chromosomal insertions of P*_c_*-*mneongreen*, P*_c_*-*mjuniper*, P*_c_*-*mscarlet-i3* and P*_c_*-*mlychee*, which represent notable advances in the *P. aeruginosa* community. The pUC18T-mini-Tn7T system allowed for stable, single-copy integration into the *P. aeruginosa* genome at high recombination rates. The selected P*_c_* promoter drove strong, stable constitutive expression of the FPs; and the four recent-generation FPs provided bright, stable and robust signals across the visible spectrum. Altogether, this design meets the needs of a wide variety of studies that require stable and robust cell labelling across generations without antibiotic selection, thereby preserving normal bacterial physiology.

Our single-cell and flow cytometry studies showed that the chromosomally tagged strains displayed brighter and more uniform signals compared to a pSEVA plasmid-based maintained with gentamicin expression. In exponentially growing *P. aeruginosa* cultures, replication events overlap, leading to several partially replicated chromosomes within individual cells; and a large fraction of the population contains more than one chromosomal equivalent, with substantial proportions containing two, three or even more chromosomal equivalents (54). Because the RK2 replication machinery maintains only about 3 plasmid copies per cell in exponential phase (39,55), a similar or even higher number of chromosome P*_c_*-*mneongreen* copies may be present in exponentially growing cells, which could reconcile the copy-number expectation with the fluorescence intensity differences between plasmid-based and chromosomally integrated construct in our experiments. Opposite results might have been observed if a high-copy plasmid had been used, yet this would have increased the risk of fitness alteration due to the fitness cost of high-copy plasmids. In addition, plasmid carrying strains were cultivated the in presence of gentamicin to ensure plasmid maintenance. Gentamicin resistance gene carried by the plasmid inactivates the gentamicin molecule and prevents its action towards ribosomes. Even if this gene is activated, residual antibiotic activity could remain, thereby exposing bacteria to sublethal antibiotic concentration, which has been shown to impact bacterial fitness (56,57). This residual effect could slow down the translation of highly expressed proteins like mNeonGreen in our case without affecting general cell fitness, thereby leading to a lower measured fluorescence intensity.

Also, cell-to-cell signal heterogeneity was greater in the plasmid-based construct than in the chromosomally inserted construct, due to fluctuations in plasmid copy number or segregation within the cell. Overall, we show that the chromosome-inserted construct outperformed the plasmid-based one and was better suited for stable expression and robust labelling in every aspect. Chromosomal insertion in Gram-negative bacteria such as *P. aeruginosa* requires a multistep engineering workflow involving transformation, successive selection steps and FLP-FRT-mediated excision of the antibiotic resistance cassette. We showed that this last excision step of mini-Tn7 insertion can be skipped, as it is not necessary for the construct functionality and the presence of the resistance-cassette had no detectable effect on bacterial fitness. Consistent with this, chromosome insertion using the pUC18T-mini-Tn7T system has been used in *P. aeruginosa* without excision of the resistance-cassette (25).

The codon-optimised mLychee and mScarlet-I3 displayed similar brightness and photostability in *P. aeruginosa* (Fig. 2D). This was unexpected, as mScarlet-I3 protein fusions have been reported to be brighter and less photostable than mLychee in *E. coli*(12). Overall, we were unable to detect fundamental differences in brightness and photostability between the two proteins. These observations might be due to high constitutive expression of the FPs compared with FP-fusion-based measurements previously described(12), and/or to differences in imaging setup, laser power, cellular environment or other factors that are not apparent under our experimental conditions.

None of the four strains carrying chromosomally inserted constructs showed altered growth at either the single-cell or the biofilm level, indicating that the fitness cost of insertion is very low or absent under the conditions tested. Each strain allowed for precise quantification of basic biofilm characteristics and allowed for monitoring biofilm development dynamics overtime accurately. Beyond structural characterization, this toolbox provides a robust platform for quantitative studies of spatial organization and population dynamics in mixed biofilms, including competition between mutants or interactions within multispecies communities. Furthermore, bronchial epithelial cell infection revealed comparable infection dynamics across the four strains, with consistent bacterial growth at each time point, indicating that virulence behaviour was not detectably affected by one specific labelling.

Interestingly, our tools revealed novel mixing dynamics in swarming colonies at the macroscopic scale. During expansion, tendrils avoided other groups of swarming cells, an avoidance known to be caused by rhamnolipids sensing (46,58), where rhamnolipids (or associated secondary metabolites) between tendrils act as a self-generated repellent that keeps tendrils separated and maximises surface coverage (51). By positioning colonies in close proximity, our experimental setup bypassed the rhamnolipid-mediated avoidance response and enabled interactions between two genetically similar tendrils, revealing an unexpected retrograde migration upon their merge. These observations also connect with the broader field of kin recognition, where bacterial populations discriminate between self and non-self and either merge or establish physical barriers depending on their relatedness. This phenomenon has been investigated in swarming bacteria such as *Bacillus subtilis* (59,60), and *Pseudomonas aeruginosa* (61). Overall, the designed fluorescent constructs allowed for robust subpopulation disclination, thus providing a powerful tool to investigate spatiotemporal population dynamics at the colony scale and uncover collective behaviours during swarming.

Taken together, our results indicate that the constructs are well suited for multiscale imaging of both live and fixed samples. Their use can be extended to multispecies co-culture imaging across imaging scales(3,6,31,62–64), biofilm dispersion studies(65,66), and evolutionary-biology experiments (23,67,68). The mini-Tn7 system has previously been used in experimental evolution studies (66) to monitor bacterial competition, thus highlighting its suitability for stable, long-term labelling during multiple generations. In addition, these constructs are also ideally suited for flow cytometry and fluorescence-activated cell sorting (FACS), allowing robust quantification of population composition, and the isolation of labelled cells for downstream genomic, transcriptomic, or physiological analyses. Lastly, our tools are particularly well-suited to challenging experimental conditions, as they are robust and stable in the absence selection pressure: Mini-Tn7 insertion for cell labelling has already been used in AirGels (25) or hydrogel (69) models for infection and biofilm studies.

Because the constructs are chromosomally integrated, they can also be combined with other genetic constructs without the risk of plasmid incompatibilities. Fluorescent transcriptional reporters or protein fusions can be carried on the chromosome or on a plasmid, allowing quantification of gene expression or precise protein localisation in complex samples(70). We have described the constructs in *P. aeruginosa* PAO1, but the pUC18T based delivery plasmids are modular and can be easily adapted. The promoter and coding sequence can be swapped to insert any desired reporter in single copy into the chromosome of *P. aeruginosa*. Finally, studies involving these constructs are not limited to *P. aeruginosa*, the pUC18T-mini-Tn7T systems is functional in *Pseudomonas putida*, *Burkholderia spp*., and *Yersinia pestis*, and likely many other Gram-negative bacteria, as most of them carry the attTn7 site (26).

In summary, by combining spectral versatility, chromosomal stability, and modular design, this toolkit provides a broadly applicable resource for imaging *P. aeruginosa* across scales, from single cells to communities and host-infection models, and offers a template that is readily transferable to other Gram-negative bacteria. We anticipate that our strains will facilitate studies of *P. aeruginosa* cell biology, community behaviour, evolution and pathogenesis.

## MATERIAL AND METHODS

### Bacterial strains and culture conditions

All mutants were created in the *P. aeruginosa* strain PAO1 background (71). Bacteria were grown in Luria-Bertani (LB) broth under aerated conditions at 160 rpm or on LB agar plates. pSEVA carrying strains are cultured in media containing 50 µg/mL gentamicin (Gm) for plasmid maintenance.

### DNA manipulations

Enzymes were used following standard molecular biology procedures. PCR reactions were performed using Q5 high-fidelity DNA polymerase (New England Biolab) or OneTaq DNA Polymerase (New England Biolabs). Restriction enzymes and T4 DNA ligase were purchased from New England Biolabs. DNA extraction was performed using QIAquick Gel Extraction Kit (QIAGEN) or QIAprep Spin Miniprep Kit (QIAGEN).

### Plasmid construction

The pUC18T-mini-Tn7T-Gm-P*_c_*-mNeonGreenPA, pUC18T-mini-Tn7T-Gm-P*_c_*-mScarlet-I3PA, pUC18T-mini-Tn7T-Gm-P*_c_*-mLycheePA, and pUC18T-mini-Tn7T-Gm-P*_c_*-mJuniperPA suicide vectors were created. Codon optimised sequences of the four FPs (12,33,34) were synthesized and sequenced by GeneCust (France) and were inserted into the pUC18T-mini-Tn7T-Gm-P*_c_* mNeonGreen(35) backbone.

Overlap extension PCR was performed using PcpromOEjoinP3 and anypromOEjoinP4 (containing StuI site) primers on the synthetised fragments SacI-rsmZ-RBS-mNeonGreen-Term-StuI and on SacI-rsmY-RBS-mScarletI3-Term-StuI amplifying the RBS-FP sequence. Meanwhile, PCR amplification using PcPromF (containing SacI site) and PcpromOEjoinP2 was performed on pUC18T-mini-Tn7T-Gm-P*_c_*-mNeonGreen to amplify the P*_c_* promoter sequence. Gel extraction of the resulting fragments was performed and one last PCR using PcPromF -anypromOEjoinP4 was done using both resulting fragments as template to join the P*_c_* sequence to the wanted FP, thus creating the two fragments SacI-P*_c_*-RBS-mNeonGreenPA-StuI and SacI-P*_c_*-RBS-mScarlet-I3PA-StuI. The fragments were gel purified for insertion in the pUC18T-mini-Tn7T-Gm backbone; a SacI/StuI double digest was performed on the pUC18T-mini-Tn7T-Gm-P*_c_*-mNeonGreen plasmid to remove the existing P*_c_*-*mneongreen* insert, as it was not codon-optimised for *P. aeruginosa*. The digested vector and fragments were joined using T4 DNA ligase resulting in the two final plasmids pUC18T-mini-Tn7T-Gm-P*_c_*-mNeonGreenPA, pUC18T-mini-Tn7T-Gm-P*_c_*-mScarlet-I3PA.

The pUC18T-mini-Tn7T-Gm-P*_c_*-mLycheePA and pUC18T-mini-Tn7T-Gm-P*_c_*-mJuniperPA plasmids were created using KpnI-RBS-mLycheePA-SphI, KpnI-RBS-mJuniperPA-SphI fragments and pUC18T-mini-Tn7T-Gm-P*_c_*-mNeonGreenPA as the vector. KpnI/SphI double digestion was performed on both fragments and the vector. These were ligated together resulting in the pUC18T-mini-Tn7T-Gm-P*_c_*-mJuniperPA and pUC18T-mini-Tn7T-Gm-P*_c_*-mLycheePA plasmids. pSEVA-Gm-P*_c_*-mNeonGreenPA was created using the pSEVA627M and pUC18T-mini-Tn7T-Gm-P*_c_*-mNeonGreenPA as matrix by Gibson cloning. The 5 plasmids were sequenced by Eurofins Genomics (France) prior to transformation into *P. aeruginosa*. pUC18T-mini-Tn7T-Gm-P*_c_*-mNeonGreenPA, pUC18T-mini-Tn7T-Gm-P*_c_*-mScarlet-I3PA, pUC18T-mini-Tn7T-Gm-P*_c_*-mLycheePA, pUC18T-mini-Tn7T-Gm-P*_c_*-mJuniperPA and pSEVA-Gm-P*_c_*-mNeonGreen plasmids were deposited at Addgene with catalog ID nos. 256432, 256433, 256434, 256435 and 256436 respectively.

### Plasmid transformation and selection

The pUC18T-mini-Tn7T based plasmid and the pTNS2 helper plasmid were transformed separately into the *E. coli* RHO3 (72) conjugative strain and conjugated via triparental mating into *P. aeruginosa* PAO1. The selection procedures were conducted as described in Choi and Schweizer(29); positive clones were tested by PCR using the PTn7R/P*glmS*-down primers. Flp-FRT excision of the gentamicin cassette was done using the pFLP3 plasmid. Gentamicin sensitive clones were tested by PCR to confirm excision of the Gm^R^ gene using Exci-F/Exci-R primers.

Antibiotics were used at the following concentrations: 100 µg/mL ampicillin for *E. coli* and 100 µg/mL gentamicin and 200 µg/mL carbenicillin for *P. aeruginosa*. 200 µg/mL diaminopimelic acid (DAP) was used to support growth of *E. coli* RHO3.

### Growth assay

Overnight cultures were diluted to an OD_600_ of 0.05 in a 96 well plate (Greiner Bio-One, France) using 200 µL of fresh LB medium. The plate was then cultured at 37 °C under agitation, and a Biotek plate reader (Agilent, USA) was used to monitor OD_600_ every 5 min for 24 h. Doubling time was determined from the growth curves of each strain.

### Single-cell imaging

Overnight cultures were diluted to an OD_600_ of 0.05 into 5 mL of fresh LB medium and incubated at 37 °C with agitation until reaching an OD_600_ of 0.1–0.2 (mid exponential phase). Then, 3 μL of culture were placed on a 1 % agarose LB pad. For the pSEVA carrying strain, both the liquid growth medium and agarose pad were supplemented with 50 µg/mL gentamicin to ensure plasmid maintenance.

Epifluorescence imaging was performed on a Zeiss Elyra PS1 automated inverted microscope with an incubation chamber (temperature set to 37 °C) and a 100 ×/1.46 NA Apochromat oil immersion objective. FP excitation was achieved using the 405 nm, 488 nm and 561 nm laser lines at a power level ranging from 1% to 20 % of the maximum output power (100 mW). The collected emission signals were subsequently filtered using a bandpass filter set (emission filters at, 420 - 480 nm, 500 - 575 nm and 570 - 650 nm for the 405 nm, 488 nm and 561 nm laser lines, respectively). Images were captured using an Andor iXon 897 EM-CCD camera with a final pixel size of 64 nm (an additional optovar lens 2.5x was placed before the camera). The camera and microscope were controlled by the ZEN software (ZEN 2012 SP2). For time-lapse experiments, images were acquired at 5 minutes intervals over 24 cycles (2 hours total) with a camera EM gain of 50. The Definite Focus was activated to compensate for the sample’s focus drift during data acquisition.

Image analysis was done using Fiji/ImageJ, fluorescence intensity was determined by measuring the mean intensity of each individual cells in 2 microcolonies at every timepoint in 3 different fields of view. Fluorescence intensities were normalised by maximum and plotted. For both P*_c_*-*mneongreen* constructs, mean intensities at T0 and T120 were normalised to background intensity and plotted. All statistical analyses were performed with GraphPad Prism 10 (GraphPad Software, LLC). Pairwise comparison between two conditions was done with a two-sided nonparametric Mann–Whitney test, *P*-values are displayed as follows: ****, *P* < 0.0001; ***, 0.0001 < *P* < 0.001; **, 0.001 < *P* < 0.01; *, 0.01 < *P* < 0.05; ns, *P* > 0.05.

### Flow cytometry

Overnight cultures were diluted to an OD_600_ of 0.05 in 5mL of fresh LB medium and grown until OD_600_ = 0.1–0.2; 5 µL of culture were diluted in 200 µL PBS. Samples were observed by Flow cytometry (Beckman Coulter’s benchtop CytoFLEX flow cytometer) recording 10,000 events per sample. A gate was previously designed based on B525-A and Y610-A signals to remove debris. Graphs were generated using the CytExpert software (Beckman Coulter, USA).

### Biofilm formation

Fluorescently labelled strains were grown overnight in LB medium supplemented with 50 µg/mL Gm to maintain plasmid in the pSEVA-carrying strains.

200 μL mono- or mixed-cultures (1/3 ratio per strain) of *P. aeruginosa* were prepared (OD_600_ = 0.05) in polystyrene 96-well microtiter plates with a μclear® base (Greiner Bio-One, France) optimised for high-resolution fluorescence imaging. After 2 h at 37 °C, the media (LB) was refreshed to discard non-adherent bacteria. Then, the microtiter plate was incubated for 4-, 12-, or 24-h at 37 °C and imaged using a Ferment du Futur Leica MICA Laser Scanning Confocal Microscopy Microscope (CLSM) at the MIMA2 INRAE imaging facility. Exposure time and laser intensity for each fluorophore was calibrated on a 24 h biofilm sample using the auto-exposure and relight parameters. When needed, biofilms were labelled with propidium iodide (PI) with a final concentration of 1 µM prior to imaging.

### Epithelial cell culture

BEAS-2B (ATCC CRL-3588) bronchial epithelial cells were cultured in Dulbecco’s Modified Eagle’s Medium (DMEM) supplemented with 10 % calf foetal serum in 75 cm^2^ cell culture flasks incubated at 37 °C, 5 % CO_2_. 48 h prior to infection, cells were passaged, enumerated and 5 000 cells were seeded onto polystyrene 96-well microtiter plates with a μclear® base (Greiner Bio-one, France) and cultured in 200 µL DMEM medium.

### Cell infection and fixation

For fixed samples imaging, DMEM was removed and replaced with 100 µL of PBS. Cells were infected with each *P. aeruginosa* strain at a MOI of 15 and incubated at 37 °C and 5 % CO_2_ for 2-, 4- or 8-h. Samples were fixed by adding 50 µL of fixation solution (PFA 2.5 %), followed by a 30 min incubation at room temperature. Samples were labelled using either 1 µM DAPI or 3.75 µM Draq5. Images were acquired using CLSM.

### Epithelial cells and biofilm CLSM imaging

Three-dimensional images were acquired on the Ferment du Futur Leica MICA widefocal microscope equipped with a HC PL APO CS2 63×/1.20 WATER objective (NA = 1.2) at the INRAE MIMA2 Imaging facility. Image stacks were collected in z-wide mode with a z-step of 1 µm (20 steps for cell infection samples, 60 steps for biofilms samples). The lateral pixel size was 0.186 µm in both the X and Y directions, with a pixel dwell time of 0.61 µs. The pinhole was set to 1 Airy unit. The fluorophores mNeonGreen, mLychee, mJuniper and mScarlet-I3 were excited sequentially using their respective laser lines, emission was collected using the fluosync spectral technology that takes in account the precise spectrum of each defined fluorophore. Three-dimensional projections of biofilm structures and epithelial cell samples were reconstructed using IMARIS 9.3.1 software (Bitplane, Switzerland). Biofilm thickness, biovolume and roughness were quantified using the BiofilmQ software. Statistical analyses were performed using GraphPad Prism 10 (GraphPad Software). Differences between groups were analysed by two-way analysis of variance (two-way ANOVA), followed by Tukey’s multiple comparisons test. ****, P < 0.0001; ***, 0.0001 < P < 0.001; **, 0.001 < P < 0.01; *, 0.01 < P < 0.05; ns, P > 0.05.

### Live cell infection and imaging

For live infection time lapses, BEAS-2B epithelial cells were labelled with a 200 nM Mito tracker Deep Red FM (Thermo Fisher Scientific) solution diluted in PBS and incubated for 30 min, 5 % CO_2_. Cells were washed with PBS, infected using a MOI of 15 and imaged directly for 10 h at 37 °C.

Time-lapse imaging was performed on a Leica MICA widefocal microscope using oblique illumination mode (IMC) for bright-field contrast and widefield epifluorescence with a resolution of 0.280 µm in X and Y and 0,508 µm in Z using 470 nm and 625 nm LED excitation. Images were acquired every 20 min during 10 h using the same illumination and exposure settings throughout the experiment. A temperature of 37 °C and 60 % humidity was maintained using the integrated OKOLAB incubation system. Image sequences were analysed using LAS X (Leica Microsystems) and Fiji/ImageJ.

### Swarming assay and imaging

94 mm diameter petri dish (Greiner Bio-one, France) containing 8 g/L nutrient broth, 0,5 % glucose, 0,4 % agar, 2,5 mM CaCl_2_. Plates were poured dried 30 min on a flat surface, and left 16 h to dry upside down. 2 µL overnight cultures were inoculated on the plate and incubated upside down at 30 °C up to 24 h.

Fluorescence time-lapse for the swarming colony experiment were captured every 5 minutes using a custom-made imaging device installed within a microbiological incubator (Heratherm IGS 100, ThermoFisher Scientific). Excitation was achieved using an LED light source (pE-4000, Coolled, UK). Image acquisition was performed using a CMOS camera (Cellcam Centro, Cairn Research, UK) with a macro lens (Navitar MVL7000, Thorlabs, USA), and a dual-band emission filter (59010m, Chroma, USA). The entire setup was controlled using μManager (http://www.micro-manager.org). Images were analysed using Fiji/ImageJ.

## Acknowledgments

We thank Armand Lablaine for valuable scientific discussions, Cyrille Billaudeau for advice in image analysis, along all members of the ProCeD and B3D lab, for helpful discussions. We are grateful to Julien Descamps, Marie-Françoise Noirot-Gros and Vlad Costache for experimental, imaging advice and training. We thank the MIMA2 facility (https://doi.org/10.15454/1.5572348210007727E12) and Ferments du Futur for CLSM observations.

## Funding

Agence Nationale de la Recherche, TARGETS, ANR-22-CE44-0016 (R.C.-L.)

European Research Council (ERC) under the Horizon2020 research and innovation program, ERC CoG No 772178 (R.C.-L.)

MICROBES 2024 Mobility program from Université Paris-Saclay (M.G.)

Atouts Pass Monde mobility grant from the Région Normandie (France) (M.G.)

## Data, code, and materials availability

Research data will be deposited on Zenodo after publication and are available from the corresponding authors upon request. Constructs can be available from the corresponding authors under a material transfer agreement. Relevant identifiers are provided in the manuscript. pUC18T-mini-Tn7T-Gm-P*_c_*-mNeonGreenPA, pUC18T-mini-Tn7T-Gm-P*_c_*-mScarlet-I3PA, pUC18T-mini-Tn7T-Gm-P*_c_*-mLycheePA, pUC18T-mini-Tn7T-Gm-P*_c_*-mJuniperPA and pSEVA-Gm-P*_c_*-mNeonGreen plasmids were deposited at Addgene with catalog ID nos. 256432, 256433, 256434, 256435 and 256436 respectively.

